# Irisin stimulates the release of CXCL1 from differentiating human subcutaneous and deep-neck derived adipocytes via upregulation of NFκB pathway

**DOI:** 10.1101/2021.07.06.451152

**Authors:** Abhirup Shaw, Beáta B Tóth, Róbert Király, Rini Arianti, István Csomós, Szilárd Póliska, Attila Vámos, Ilma R Korponay-Szabó, Zsolt Bacso, Ferenc Győry, László Fésüs, Endre Kristóf

**Author notes:** Authors to whom correspondence should be addressed, **Correspondence:** László Fésüs, Endre Kristóf. These authors have contributed equally to this work and share first authorship (Abhirup Shaw, Beáta B Tóth). These authors have contributed equally to this work and share last authorship (László Fésüs, Endre Kristóf).

## Abstract

Thermogenic brown and beige adipocytes play an important role in combating obesity. Recent studies in rodents and humans have indicated that these adipocytes release cytokines, termed “batokines”. Irisin was discovered as a polypeptide regulator of beige adipocytes released by myocytes, primarily during exercise. We performed global RNA sequencing on adipocytes derived from human subcutaneous and deep-neck precursors, which were differentiated in the presence or absence of irisin. Irisin did not exert an effect on the expression of characteristic thermogenic genes, while upregulated genes belonging to various cytokine signaling pathways. Out of the several upregulated cytokines, *CXCL1*, the highest upregulated, was released throughout the entire differentiation period, and predominantly by differentiated adipocytes. Deep-neck area tissue biopsies also showed a significant release of CXCL1 during 24 hours irisin treatment. Gene expression data indicated upregulation of the NFкB pathway upon irisin treatment, which was validated by an increase of p50 and decrease of IкBα protein level, respectively. Continuous blocking of the NFκB pathway, using a cell permeable inhibitor of NFκB nuclear translocation, significantly reduced CXCL1 release. The released CXCL1 exerted a positive effect on the adhesion of endothelial cells. Together, our findings demonstrate that irisin stimulates the release of a novel “batokine”, CXCL1, via upregulation of NFκB pathway in neck area derived adipocytes, which might play an important role in improving tissue vascularization.

## 1 Introduction

Recent studies indicated the presence of thermogenic adipose tissue, capable of dissipating energy as heat under sub-thermal conditions in healthy human adults (1), (2). These are located in cervical, supraclavicular, axillary, mediastinal, paravertebral, and abdominal depots (3), (4), (5); supraclavicular, deep-neck (DN), and paravertebral having the highest amounts. Together these depots account for 5% of basal metabolic rate in adults, highlighting their importance in combating obesity and type 2 diabetes mellitus (6). In rodents, these thermogenic adipocytes are either classical brown or beige depending on their origin and distribution (7), (8). In addition to their role in thermogenesis, these adipocytes also secrete adipokines, termed ‘batokines’, which have been shown to exert autocrine, paracrine, or endocrine activity (9). For example, vascular endothelial growth factor A (VEGF-A) secreted by brown adipocytes promotes angiogenesis and vascularization of brown adipose tissue (BAT) (10), (11), (12) while Fibroblast growth factor (FGF) 21 enhances the beiging of white adipose tissue (WAT) and increases thermogenesis in BAT (13), (14), (15). Understanding the roles of batokines in the human body is an area of active research (16), (17).

Irisin, a cleaved product of the transmembrane protein FNDC5, was discovered as a myokine in mice and was shown to be a browning inducing endocrine hormone (18), (19), presumably acting via integrin receptors (20). In mice, irisin secretion was induced by physical exercise and shivering of skeletal myocytes, which induced a beige differentiation program in subcutaneous WAT (19). In humans, inconsistent effects were found when adipocytes of different anatomical origins were treated with recombinant irisin (21), (22), (23), (24), (25), (26). How irisin affects the differentiation of the thermogenically prone neck area adipocytes still awaits description. We have previously reported that human DN adipose tissue biopsies released significantly higher amounts of interleukin (IL)-6, IL-8, monocyte chemoattractant protein 1 (MCP1) as compared to subcutaneous ones, which was further enhanced upon irisin treatment (27). CXCL1, previously known as growth-related oncogene (GRO)-α, is a small peptide belonging to the CXC chemokine family; newly synthetized CXCL1 by vessel-associated endothelial cells and pericytes facilitates the process of neutrophil diapedesis (28).

In this study, we aimed to get an overview of all the genes whose expression is regulated by irisin. For this, we have performed a global RNA-Sequencing comprising of *ex vivo* differentiated adipocytes of subcutaneous and deep depots of human neck from 9 individuals and analysed the upregulated genes upon irisin treatment. Surprisingly, several genes which encode secreted proteins were upregulated. Out of those, chemokine C-X-C motif ligand (CXCL) 1 was found to be the highest expressed and a novel batokine induced in differentiating adipocytes of both origins. The CXCL1 release was stimulated predominantly via the upregulation of nuclear factor-κB (NFκB) pathway. We found that the secreted CXCL1 had an adhesion promoting effect on endothelial cells, supporting that irisin can exert effects not directly linked to heat production.

## 2 Materials and methods

### 2.1 Materials

All chemicals were obtained from Sigma Aldrich (Munich, Germany) unless otherwise stated.

### 2.2 Isolation, cell culture, differentiation, and treatment of hASCs

Human adipose-derived stromal cells (hASCs) were obtained from stromal-vascular fractions of subcutaneous neck (SC) and DN tissues of volunteers, aged between 20-65 years, undergoing planned surgical treatment. A pair of biopsies from SC and DN areas was obtained from the same donor, to avoid inter-individual variations. Patients with known diabetes, malignant tumour or with abnormal thyroid hormone levels were excluded from the study. Written informed consent was obtained from all participants before the surgery.

hASCs were isolated and cultivated as previously described (27), (29). The absence of mycoplasma was confirmed by PCR analysis (PCR Mycoplasma Test Kit I/C, Promocell, Heidelberg, Germany). Cells were differentiated following a reported white adipogenic differentiation protocol, with or without the addition of human recombinant irisin (Cayman Chemicals, MI, USA) at 250 ng/mL concentration (27), (30). Media were changed every other four days and cells were used after 14 days of differentiation. Where indicated, cells were treated with RGDS peptide (10 μg/mL, R&D systems, MN, USA) (20) or SN50 (50 μg/mL, Med Chem Express, NJ, USA) (31).

### 2.3 RNA isolation, RT-qPCR, and RNA-Sequencing

Cells were collected in Trizol reagent (Thermo Fisher Scientific, MA, USA) and RNA was isolated manually by chloroform extraction and isopropanol precipitation. To obtain global transcriptome data, high throughput mRNA sequencing was performed on Illumina Sequencing platform (29). Grouping was performed based on Panther Reactome pathways (https://pantherdb.org). Heatmap visualization was performed on the Morpheus web tool (https://software.broadinstitute.org/morpheus) using Pearson correlation of rows and complete linkage based on calculated z-score of DESeq normalized data after log_2_ transformation (29). The interaction networks were determined using STRING (https://string-db.org) and constructed using Gephi 0.9.2 (https://gephi.org). The size of the nodes was determined based on fold change (29).

For RT-PCR, RNA quality was evaluated by spectrophotometry and cDNA was generated by TaqMan reverse transcription reagents kit (Thermo Fisher Scientific) followed by qPCR analysis (32).

### 2.4 Antibodies and Immunoblotting

Samples were collected, separated by SDS-PAGE, and transferred to PVDF Immobilon-P transfer membrane (Merck-Millipore, Darmstadt, Germany) as previously described (32). The following primary antibodies were used overnight in 1% skimmed milk solution: anti-p50 (1:1000, 13755, Cayman Chemicals), anti-IκBα (1:1000, 4812, Cell Signaling Technology, MA, USA), and anti-ß-actin (1:5000, A2066, Novus Biologicals, CO, USA). HRP-conjugated goat anti-rabbit (1:10,000, Advansta, CA, USA, R-05072-500) or anti-mouse (1:5000, Advansta, R-05071-500) IgG were used as secondary antibodies, respectively. Immobilion western chemiluminescence substrate (Merck-Millipore) was used to visualize the immunoreactive proteins. FIJI was used for densitometry.

### 2.5 Immunostaining analysis and image analysis

hASCs from SC and DN areas were plated and differentiated in 8 well Ibidi μ-chambers (Ibidi GmbH, Gräfelfing, Germany). Cells were treated with Brefeldin A (100 ng/mL) 24 hours prior collection to sequester the released CXCL1 (27), (31). After that, cells were washed with PBS, fixed by 4% paraformaldehyde, permeabilized with 0.1% saponin and blocked by 5% milk as per described protocols (32). The cells were incubated subsequently with anti-CXCL1 primary antibody (1:100, 712317, Thermo Fisher Scientific) and Alexa 488 goat anti-rabbit IgG (1:1000, A11034, Thermo Fischer Scientific) secondary antibody for 12 and 3 hours at room temperature, respectively. Propidium iodide (1.5 μg/mL, 1 hour) was used to label the nuclei. Images were acquired with Olympus FluoView 1000 confocal microscope and analysed by FIJI as described previously (32). Adipogenic differentiation rate was quantified as described previously (25), (33).

### 2.6 Determination of the released factors

Supernatants of samples from cell culture experiments were collected at the regular replacement of the media, on days 4, 12, 18, 21 of differentiation, wherever indicated. For SC and DN, supernatants were collected and stored at −20 oC from the differentiated cells of the same donor and considered as one repetition, followed by repetition with subsequent donors. For tissues, 10–20□ mg of SC and DN tissue samples from the same donor were floated for 24□ hours in DMEM-F12-HAM medium with or without the presence of 250 ng/mL irisin (27), (34). The release of CXCL1, CX3CL1, IL-32, TNFα and IL1-β were analysed from the stored samples using ELISA Kits (R&D systems, MN, USA).

### 2.7 Adhesion assay

Human Umbilical Vein Endothelial Cells (HUVEC) cell line, HUCB2 was generated from endothelial cells isolated from the human umbilical cord vein of a healthy newborn by collagenase digestion as described earlier (35). Cells were cultured in M199 medium (Biosera, Nuaille, France) containing 10% FBS (Thermo Fisher Scientific), 10% EGM2 Endothelial Growth Medium (Lonza, Basel, Switzerland), 20 mM HEPES (Biosera), 100 U/mL Penicillin, 100 μg/mL Streptomycin and 2.5 μg/mL Amphotericin B (Biosera), and immortalized by the viral delivery of telomerase gene using pBABE-neo-hTERT (36) (gift from Bob Weinberg, 1774, Addgene). The virus packaging was performed in HEK293FT cells (Thermo Fisher Scientific) based on a calcium precipitation method using pUMVC and pCMV-VSV-G vectors (37) (gift from Bob Weinberg, 8449 and 8454, Addgene). The pseudovirion containing supernatant was used for infection, and selection was started 72 hours later using 300 μg/mL G418 (Merck-Millipore). Immortalized cells completely retain the morphological properties of primary endothelial cells.

Prior to the adhesion assay, EGM2 was omitted from the standard medium of HUCB2 cells and FBS content was decreased to 1% for 24 hours. 96-well plates (Thermo Fisher Scientific) were precoated with fibronectin (Merck-Millipore) at 1.25 μg/mL concentration in PBS, for 1 hour at 37oC and then washed twice with PBS. Cells were plated at 1000 cells/well density and left to adhere for 2 hours in the CO_2_-incubator in the mixture (1:1 ratio) of starvation and conditioned media (incubation period from day 8 to 12 of differentiation) from SC and DN adipocytes, differentiated in the presence or absence of 250 ng/mL irisin, respectively. Where indicated, recombinant human CXCL1 (275-GR, R&D Systems) was used at 2500 pg/mL concentration in starvation media. Unattached cells were removed by once washing with PBS and adhered cells were incubated with starvation media containing CellTiter-Blue Cell Viability reagent (resazurin; Promega, WI, USA; 36 times dilution). To determine the ratio of attached cells in various conditions, the fluorescent intensity change of each well (Ex:530nm/Em:590nm), due to the conversion of resazurin to resorufin by cellular metabolism, was measured using Synergy H1 (BioTek, Hungary) plate reader 2, 4, 6, 18, and 24 hours after adding resazurin. The effects of conditioned media and recombinant CXCL1 on the adhesion were expressed as fold changes of the fluorescent intensity growth rate (slope) relative to their respective controls after subtraction of only starvation media and only cells from each value.

### 2.8 Statistics and Image analysis/preparation

Results are expressed as mean±SD for the number of independent repetitions indicated. For multiple comparisons of groups, statistical significance was determined by one-way analysis of variance followed by Tukey *post hoc* test. In comparison of two groups, two-tailed paired Student’s t-test was used. For the design of graphs and evaluation of statistics, Graphpad Prism 9 was used.

## 3 Results

### 3.1 Irisin did not change the differentiation potential of adipocytes while increased the expression of integrin receptor genes in both subcutaneous (SC) and deep-neck (DN) origins

Primary hASCs from 9 independent donors were isolated and cultivated from SC and DN area of human neck, as described (29). Adipogenic differentiation was driven by a white adipocyte differentiation medium with or without the presence of irisin for 14 days. Then, the global gene expression pattern of differentiated adipocytes and undifferentiated hASCs were determined by global RNA-sequencing (29). Gene expression of general adipocyte markers (e.g. *FABP4, ADIPOQ*) was higher in all differentiated adipocytes as compared to preadipocytes (Figure 1A). Quantification of the adipogenic differentiation rate by laser-scanning cytometry (25) revealed that more than 50% of the cells were differentiated following our 14-days long differentiation protocol (Figure 1B). The presence of irisin did not affect the differentiation and gene expression of general adipocyte markers (Figure 1 A,B). A recent publication proposed the irisin receptors to be integrins (ITGAV-ITGB1/3/5) (20). Hence the expression of *ITGAV* was analysed from RNA-sequencing data (Figure 1C), which revealed that it is expressed in both the preadipocytes and differentiated adipocytes. Upon RT-qPCR validation, a significant increase of *ITGAV* expression was observed in both SC and DN adipocytes in response to irisin (Figure 1D). RNA-sequencing data showed that *ITGB1, 3*, and *5* were also expressed at a high extent in preadipocytes and in differentiated adipocytes irrespective of the presence of irisin (Supplementary Figure 1).

**Figure 1.**
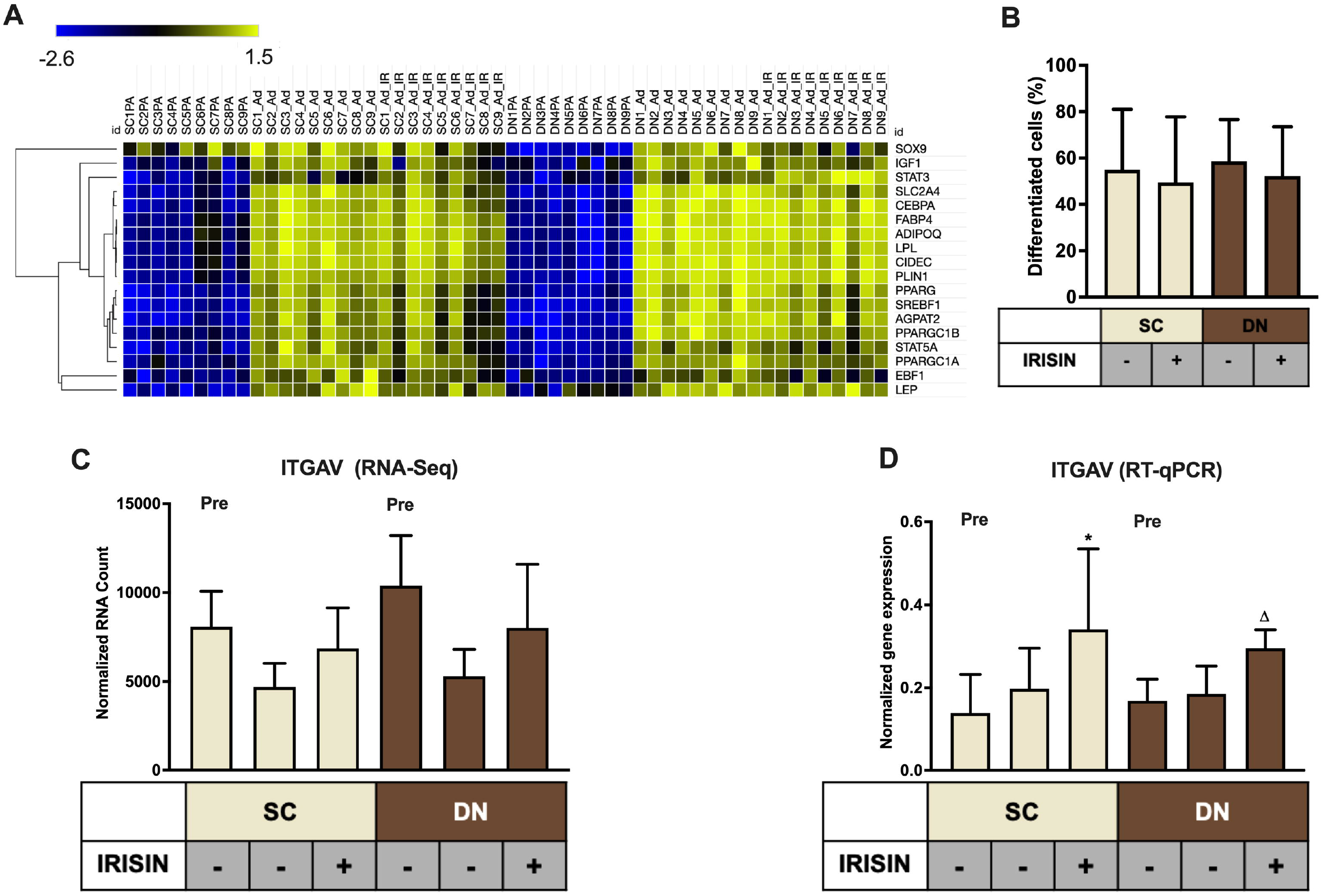
Preadipocytes from subcutaneous (SC) and deep-neck (DN) depots of human neck differentiated at a similar extent irrespective of the presence of irisin. SC and DN preadipocytes (Pre) were differentiated for two weeks to white adipocytes. Where indicated, 250 ng/ml irisin was administered during the whole differentiation process. (A) Heatmap illustrating the expression of general adipogenic differentiation markers in samples used for Global RNA Sequencing (n=9), (B) Quantification of differentiation rate by laser-scanning cytometry (n=9), (C) Quantification of *ITGAV* gene expression determined by RNA Sequencing (n=9) and (D) RT-qPCR, normalized to *GAPDH* (n=5). Data presented as Mean ± SD. *: Refers to compared with SC, Δ: Refers to compared with DN. *,^Δ^ p<0.05. Statistics: paired t-test (D).

### 3.2 Genes involved in chemokine signaling pathways were upregulated in adipocytes differentiated with irisin

RNA-Sequencing analysis identified 79 genes to be higher expressed upon irisin treatment that are visualized by a Volcano plot (Figure 2A). 50 and 66 genes were significantly upregulated in SC and DN area adipocytes, respectively, each of which are listed in Supplementary Table 1. 37 genes, including *CXCL1, CX3CL1, IL32, IL34, IL6*, and *CCL2* were found to be commonly upregulated in adipocytes of both depots (Figure 2A, 2B, Supplementary Table 1). Surprisingly, thermogenic marker genes did not appear among these. Panther enrichment analysis of genes upregulated in both SC and DN adipocytes by irisin treatment revealed pathways such as cytokine signaling (*NFKB2, CXCL1, CXCL2, IL32, IL34, IL6, CCL2*), interleukin-4 and 13 signaling (*IL6*, *CCL2, JUNB, ICAM1*), and class A/1 rhodopsin like receptors (*CXCL3*, *CXCL5, CX3CL1, CXCL2, CCL2, CXCL1*), which were commonly upregulated in both SC and DN adipocytes (Table 1). Gephi diagrams illustrate the interaction of upregulated genes belongs to several pathways (Figure 2-D). Interleukin-10 signaling were amongst the upregulated pathways in SC adipocytes (Figure 2C), while in DN, G-alpha-I and response to metal ions were upregulated (Figure 2D). Cluster analyses and heatmap illustration of the gene expression values of the 79 higher expressed genes upon irisin treatment identified two main clusters: a cluster of 25 genes that uniquely expressed in irisin treated mature adipocytes, and another group of genes that are expressed highly in preadipocytes, but suppressed in differentiated adipocytes without irisin treatment (Supplementary figure 2). The higher expression of *IL6, CCL2, CX3CL1*, and *IL32*, cytokine encoding genes was observed by both RNA Sequencing and RT-qPCR analysis (Supplementary figure 3). Release of IL-6 and MCP1, encoded by *CCL2*, was detected from conditioned media collected during differentiation and was found to be specifically released by differentiated lipid laden adipocytes as described in our previous publication (27). Next, we investigated if fractalkine (encoded by *CX3CL1* gene) and IL-32 were released into the conditioned media collected during the differentiation on days number 4 and 12; however, we were unable to detect these factors (data not shown).

**Figure 2.**
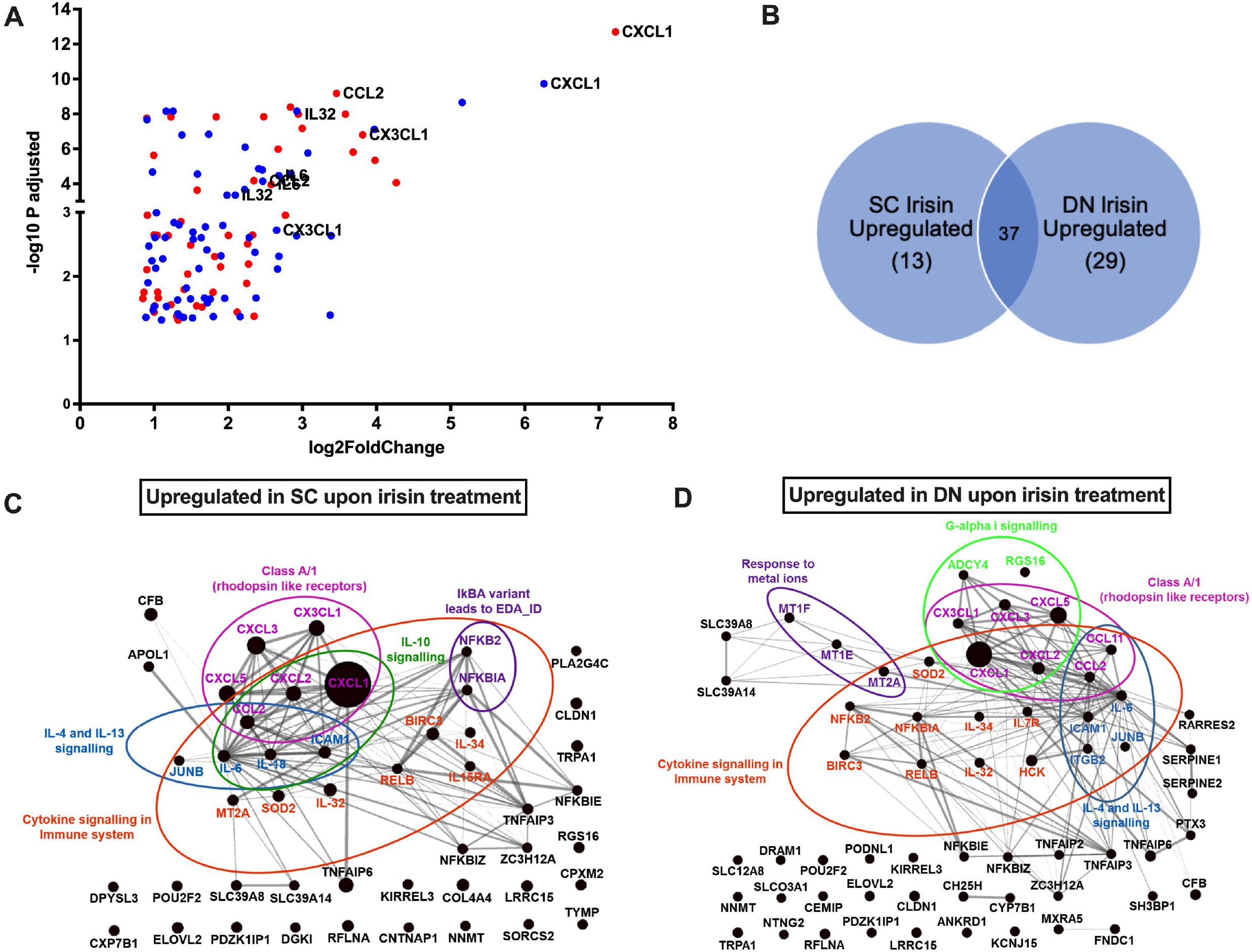
Irisin upregulated similar gene-sets that encode for cytokines in subcutaneous (SC) and deep-neck (DN) depots of human neck area adipocytes. SC and DN preadipocytes were differentiated and treated as in Figure 1. (A) Volcano plot showing each of the upregulated genes in SC (red) and DN (blue) depots upon irisin treatment; the highest upregulated genes are listed separately, (B) Venn-diagram illustrating the genes commonly upregulated by irisin treatment in SC and DN depots. Gephi illustrations highlighting the most important pathways and the interaction of genes upregulated by irisin treatment in SC (C) and DN (D) derived adipocytes.

**Table 1.** Pathways of significantly upregulated genes upon irisin treatment during differentiation of subcutaneous (SC) and deep-neck (DN) derived adipocytes. Genes commonly upregulated in both SC and DN area adipocytes are marked red. *CXCL1* was the highest upregulated gene in both SC and DN area adipocytes. FDR: False Discovery Rate.

| SC Irisin Upregulated |  |  |
| --- | --- | --- |
| Panther Reactome Pathways | Gene name | FDR |
| IkBA variant leads to EDA-ID | *NFKBIA, NFKB2* | 4.49x10<sup>-2</sup> |
| Cytokine signaling in immune system | *IL6, NFKBIA, JUNB, IL32, SOD2, MT2A, NFKB2, CXCL2, CCL2, IL15RA, IL18, IL34, ICAM1, CXCL1, RELB, BIRC3* | 5.23x10<sup>-8</sup> |
| Interleukin-10 signaling | *IL6, CXCL2, CCL2, IL18, ICAM1, CXCL1* | 1.65x10<sup>-6</sup> |
| Class A/1 (Rhodopsin like receptors) | *CXCL3, CXCL5, CX3CL1, CXCL2, CCL2, CXCL1* | 3.5x10<sup>-2</sup> |
| Interleukin-4 and Interleukin-13 signaling | *IL6, JUNB, CCL2, IL18, ICAM1* | 2.3x10<sup>-3</sup> |
| DN Irisin Upregulated |  |  |
| Panther Reactome Pathways | Gene name | FDR |
| Response to metal ions | *MT2A*, MT1E, MT1F | 4.74x10<sup>-3</sup> |
| Class A/1 (Rhodopsin like receptors) | *CCL11*, *CXCL3*, *CXCL5*, *CX3CL1*, *CXCL2*, *CCL2*, *CXCL1* | $1.85 \times 10^{-2}$ |
| Cytokine signaling in immune system | *IL6*, *CCL11*, *ITGB2*, *NFKBIA*, *JUNB*, *IL32*, *SOD2*, *MT2A*, *NFKB2*, *IL7R*, *CXCL2*, *CCL2*, *IL34*, *ICAM1*, *HCK*, *CXCL1*, *RELB*, *BIRC3* | $5.55 \times 10^{-8}$ |
| Interleukin-4 and Interleukin-13 signaling | *IL6*, *CCL11*, *ITGB2*, *JUNB*, *CCL2*, *ICAM1* | $6.33 \times 10^{-4}$ |
| G-alpha (i) signaling events | *CXCL3*, *CXCL5*, *CX3CL1*, *ADCY4*, *RGS16*, *CXCL2*, *CXCL1* | $5.07 \times 10^{-2}$ |

### 3.3 Irisin dependent induction of CXCL1 release occurred predominantly from differentiating and mature adipocytes

Irisin upregulated *CXCL1* gene expression at the largest extent in both SC and DN area adipocytes (Figure 2A, 3A, Supplementary table 1). This observation was verified by RT-qPCR (Figure 3B). As a next step, release of CXCL1 from irisin treated and untreated adipocytes was investigated into the conditioned differentiation media collected on the fourth and twelfth days of differentiation. Irisin treatment resulted in significant increase in CXCL1 secretion at the intervals of days 0-4 and 8-12 in both types of adipocytes (Figure 3C).

**Figure 3.**
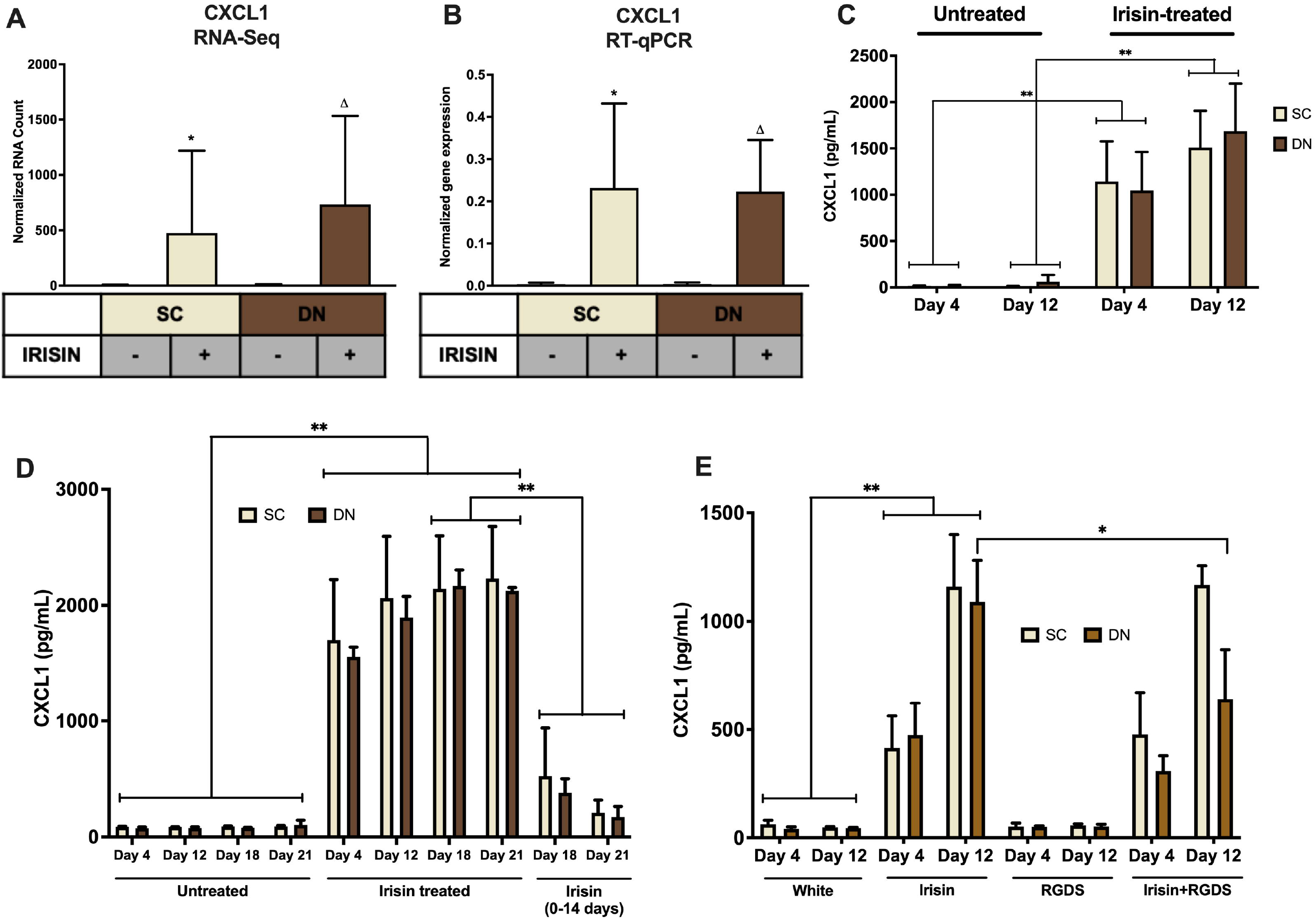
Irisin dependent CXCL1 release was stimulated from differentiating subcutaneous (SC) and deep-neck (DN) area adipocytes. SC and DN preadipocytes were differentiated and treated as in Figures 1-2. Where indicated, irisin was omitted from the differentiation medium at day 14. Conditioned differentiation media was collected and secreted CXCL1 was measured by sandwich ELISA. (A) Quantification of *CXCL1* gene expression as determined by RNA Sequencing (n=9) or RT-qPCR (B) normalized to *GAPDH* (n=5), (C) CXCL1 release by *ex vivo* differentiating SC and DN adipocytes into the conditioned media collected at the indicated intervals, in the presence or absence of irisin (n=4), (D) CXCL1 release in conditioned medium collected at indicated intervals from untreated (21 days) and irisin treated (14 and 21 days as indicated) cell-culture samples (n=3), (E) CXCL1 release from differentiating adipocytes with or without irisin treatment, in the presence or absence of 10 μg/ml RGDS (n=4). Comparisons are for the respective days in case of ELISA. Data presented as Mean ± SD. *: Refers to compared with SC, Δ: Refers to compared with DN. *,^Δ^ p<0.05, **p<0.01. Statistics: GLM (A), One-way ANOVA with Tukey’s post-test (B-E).

We aimed to further investigate the dependence of CXCL1 release on the presence of irisin. Therefore, we differentiated hASCs for 21 days, with three sets of samples, each from SC and DN derived adipocytes. Two sets of hASCs were differentiated as previously described, and for the third set, irisin treatment was discontinued after 14 days. Conditioned media were collected on days number 4, 12, 18, 21 and measured for the release of CXCL1. Large amounts of CXCL1 were secreted throughout the differentiation period in the presence of irisin; however, discontinuation of irisin administration led to gradual and significant reduction of the released chemokine (Figure 3D).

A recent publication indicated that RGDS peptide, an integrin receptor inhibitor, can potentially inhibit the effect of irisin (20). Hence, we checked the effect of this peptide on the release of CXCL1 on top of irisin treatment. RGDS partially reduced the irisin-stimulated release of CXCL1 by DN adipocytes at day 12 of the differentiation period (Figure 3E).

Release of CXCL1 throughout the whole differentiation period raised a possibility that both undifferentiated preadipocytes and differentiated adipocytes are able to release the chemokine. To investigate this, the secretion machinery of the mixed cell population was inhibited, followed by CXCL1 immunostaining and image acquisition by confocal microscopy. Irisin treatment significantly increased CXCL1 immunostaining intensity in both SC (Figure 4A) and DN adipocytes (Figure 4B). Irisin treated adipocytes accumulated significantly more CXCL1 compared to their preadipocyte counterparts in both SC (Figure 4A) and DN areas (Figure 4B). Secondary antibody control images proved the specificity of the primary antibody used (Supplementary figure 4). Our data suggests that irisin stimulates the release of CXCL1 from differentiating and mature adipocytes which is strongly dependent on the presence of irisin but not prominently on its presumed integrin receptor.

**Figure 4.**
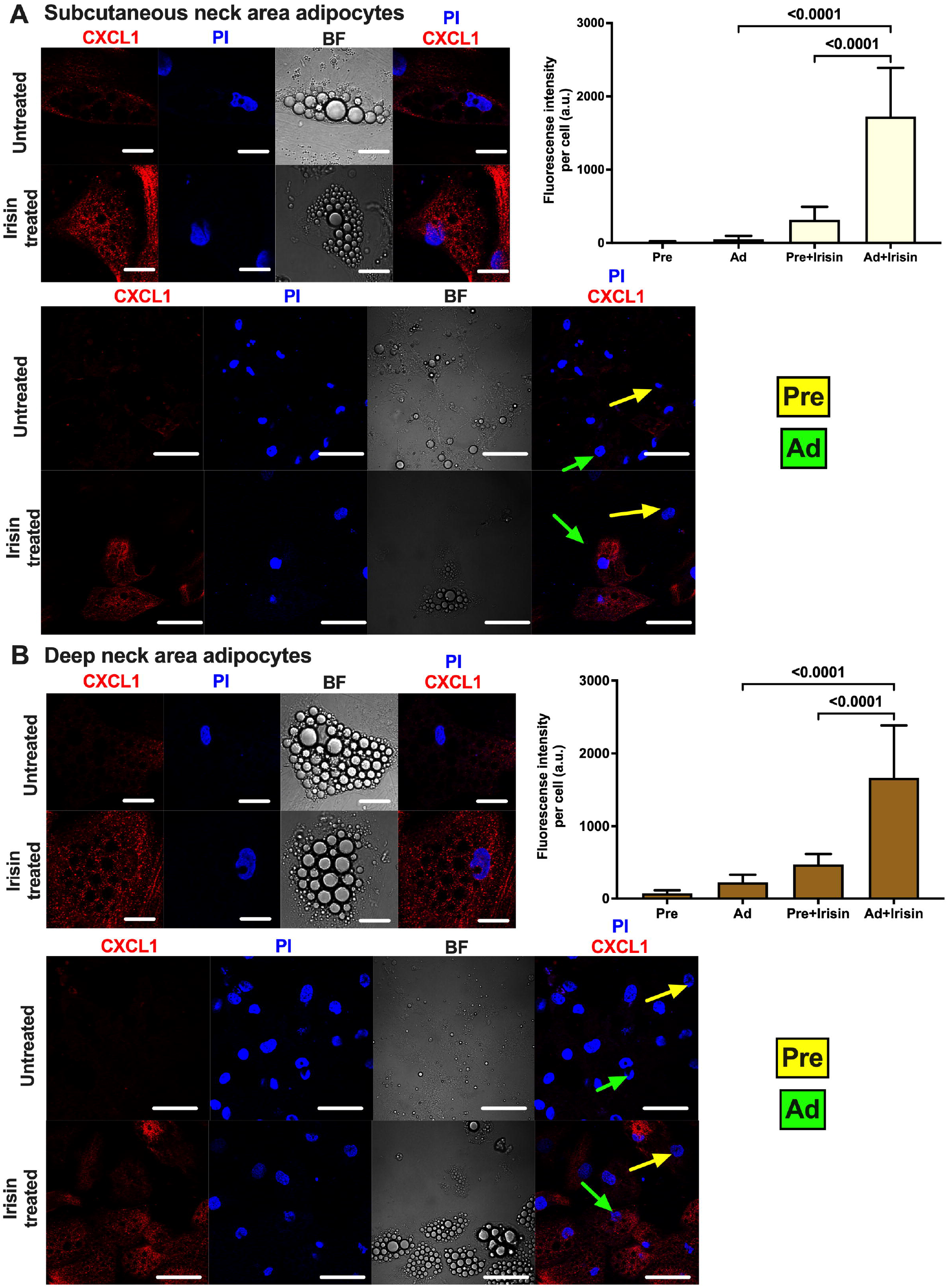
Irisin stimulated CXCL1 release predominantly from subcutaneous (A) and deep-neck (B) area differentiated adipocytes. SC and DN preadipocytes (Pre) were plated and differentiated into adipocytes (Ad) on Ibidi chambers, with or without irisin treatment as in Figures 1-3. Cells were treated with 100 ng/ml brefeldin-A for 24 hours to block the secretion of CXCL1, which was followed by fixation and image acquisition by confocal microscopy. Propidium Iodide (PI) was used to stain the nucleus. BF represents the bright field image. Confocal images of differentiated adipocytes were shown followed by wider coverage of undifferentiated and differentiated adipocytes. Scale bars represent 10 μm for single differentiated Ad and 30 μm for wider coverage of Pre and Ad. Yellow and green arrows point the undifferentiated preadipocytes and the differentiated adipocytes, respectively. Quantification of fluorescence intensity normalized to per cell are shown on the right bar graphs. Data presented as Mean ± SD. n= 35 cells (A) and 50 cells (B) from two independent donors. Statistics: One-way ANOVA with Tukey’s post-test.

### 3.4 Irisin stimulates the release of CXCL1 via the upregulation of NFκB pathway

Next, we aimed to investigate the molecular mechanisms underlying the irisin-induced CXCL1 release. According to our RNA Sequencing data, irisin treatment resulted in a significant upregulation of *NFKB2* and an increasing trend was observed for *NFKB1* and *RELA* (Supplementary figure 5A-C) genes. RT-qPCR validation indicated significant upregulation of *NFKB1* (p50 subunit) and *RELA* (p65 subunit) in DN, while an increasing trend was observed in SC adipocytes (Figure 5A-B). p50 protein expression was significantly increased in DN and an increasing trend was found in the case of SC adipocytes (Figure 5C). Protein expression of IκB.α, the inhibitor of NFκB transcription factor, decreased significantly upon irisin treatment in SC and a decreasing trend was observed in DN adipocytes (Figure 5D), indicating the upregulation of NFκB pathway.

**Figure 5.**
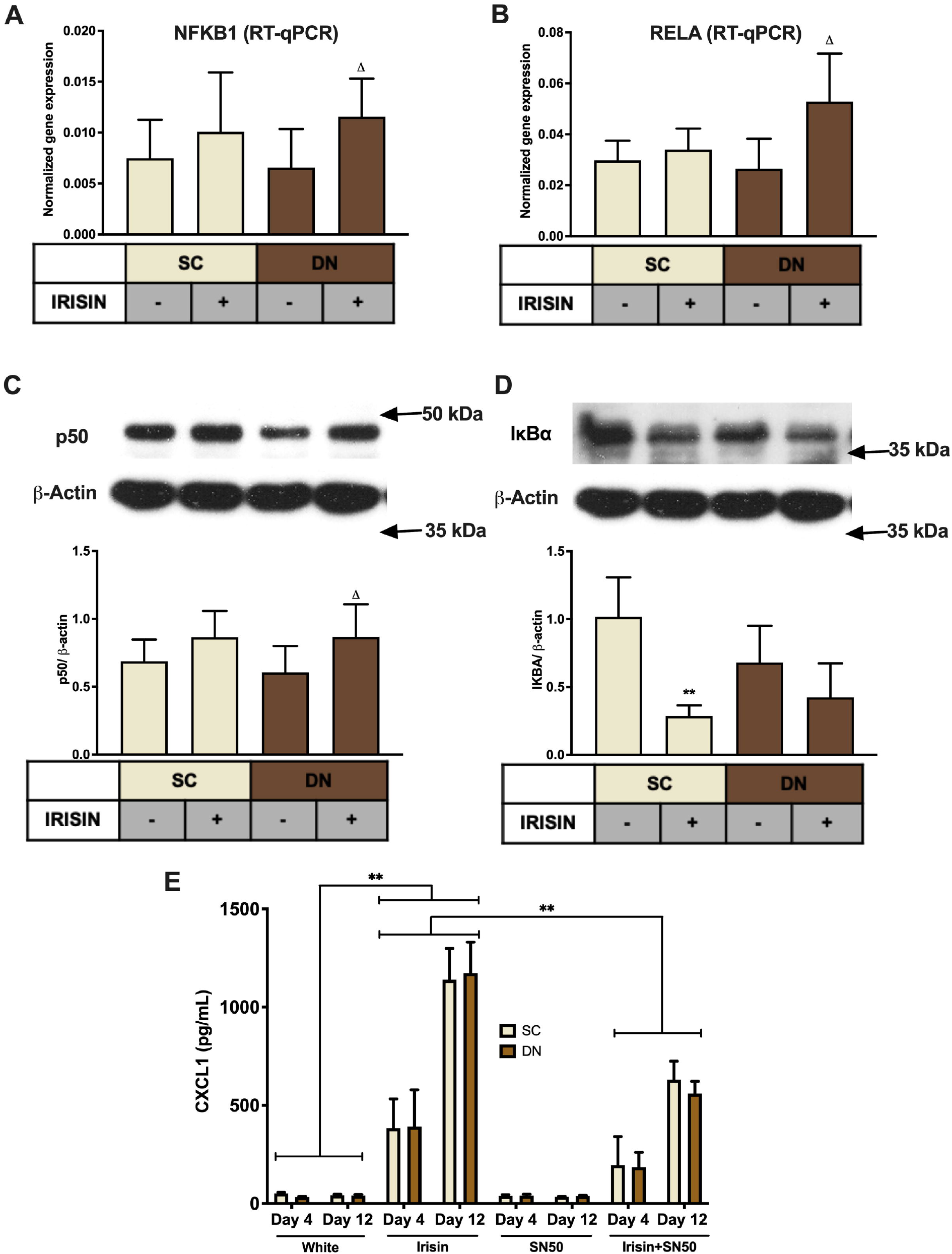
CXCL1 release is stimulated via the NFκB pathway during the differentiation of subcutaneous (SC) and deep-neck (DN) area adipocytes. SC and DN preadipocytes were differentiated and treated as in Figures 1-4. Quantification of gene expression for *NFKB1* (A) and *RELA* (B), normalized to *GAPDH* by RT-qPCR (n=5), (C) p50 and IKBA (D) protein expression, normalized to ß-actin (n=6), (E) CXCL1 release from differentiating adipocytes with or without irisin treatment, in the presence or absence of 50 μg/ml SN50 (n=4). Data presented as Mean ± SD. *: Refers to compared with SC, Δ: compared with DN, comparisons are for the respective days in case of ELISA. *^,Δ^p<0.05, **’^ΔΔ^p<0.01. Statistics: One-way ANOVA with Tukey’s post-test.

To prove the direct involvement of the NFκB pathway in adipocyte response to irisin, we applied a cell permeable inhibitor of NFκB nuclear translocation, SN50 (31), which significantly reduced the release of the chemokine from both types of adipocytes, when it was applied on top of irisin on both the fourth and twelfth days of differentiation, as compared to cells stimulated only by irisin (Figure 5E).

The observed effects of irisin are not likely to be caused by any contamination of endotoxins, which is proved by the negligible expression of *TNFα* or *CCL3* genes (Supplementary figure 5D,E), and the decreasing trend of *IL1β* gene expression (Supplementary figure 5F) in irisin treated adipocytes. Furthermore, we did not detect secreted TNFα or IL-1β in the conditioned media of either untreated or irisin treated SC and DN derived adipocytes (data not shown).

### 3.5 CXCL1 released from irisin stimulated adipocytes and adipose tissue improves the adhesion of endothelial cells

Finally, SC and DN paired tissue biopsies were floated in the presence or absence of irisin dissolved in empty media, followed by quantification of CXCL1 release. The secretion of the chemokine was significantly stimulated from DN tissue biopsies upon irisin treatment (Figure 6A).

**Figure 6.**
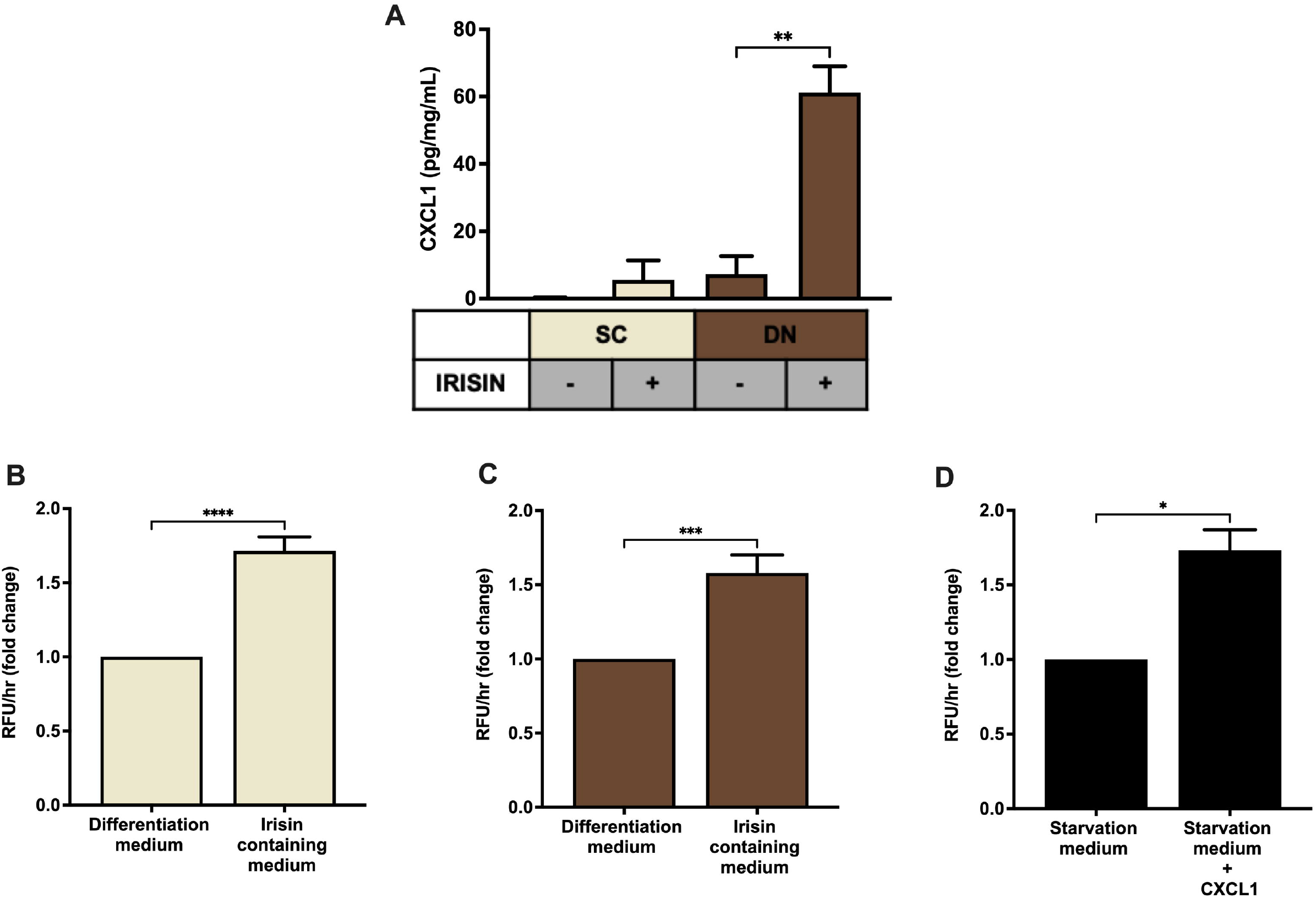
Irisin stimulated the release of CXCL1 from DN tissue biopsies, which improves the adhesion of endothelial cells. (A) CXCL1 released into the conditioned media of paired SC and DN biopsies after 24 hours incubation in the presence or absence of irisin (n=4), Quantification of adhesion of endothelial cells upon incubation with the conditioned media (with or without irisin treatment) from *ex vivo* differentiated (incubation period from day 8 to 12 of differentiation) SC (B) and DN (C) area adipocytes (n=5), (D) Quantification of endothelial cell adhesion upon incubation with recombinant CXCL1 in starvation medium (n=3). Data presented as Mean ± SD. * p<0.05, **p<0.01, ***p<0.001. Statistics: One-way ANOVA with Tukey’s post-test (A) and paired t-test (B-D).

Secretion of CXCL1 plays an important role in wound repair and angiogenesis (28). To prove whether the released CXCL1 can lead to increased adhesion of endothelial cells, conditioned media collected on the twelfth day of *ex vivo* differentiation, from untreated and irisin treated SC and DN area adipocytes, were added to HUVECs followed by a resorufin based adhesion assay. The conditioned medium from irisin treated adipocytes, which contains various released factors (including CXCL1) was able to significantly increase the adhesion of HUVECs, compared to the conditioned medium of untreated adipocytes (Figure 6B,C). When HUVECs were treated with recombinant CXCL1, at the highest observed concentration in media of irisin-treated *ex vivo* differentiated adipocytes, their adhesion was enhanced significantly (Figure 6D). This suggests a potential beneficial role of the released CXCL1 in promoting endothelial functions and adipose tissue remodelling to support efficient thermogenesis indirectly by enhancing vascularisation.

## 4 Discussion

Primarily, irisin was discovered as a proteolytic product of FNDC5, released by cardiac and skeletal myocytes, which induces a beige differentiation program in mouse subcutaneous WAT (19), (38). In humans, Adenine was replaced by Guanine in the start codon of the murine *FNDC5* gene, probably resulting in a shorter precursor protein lacking the part from which irisin is cleaved (22). Despite this, the presence of irisin in human blood plasma could be detected using mass spectrometry or different antibodies, however, in a more than 10-fold lower concentration than in rodents (39). Furthermore, it is present in the cerebrospinal fluid, liver, pancreas, stomach, saliva, and urine (40). Controversial effects were observed when differentiating human adipocytes of distinct anatomical origins were treated with the recombinant hormone (21), (22), (23), (24), (25), (26), (41). We reported that irisin induced a beige phenotype of human primary abdominal subcutaneous and Simpson-Golabi-Behmel syndrome (SGBS) adipocytes when they were treated at a concentration detected in physically active rodents on top the white adipogenic protocol that was used in this study (26), (25). Irisin administration also facilitated the secretion of batokines, such as IL-6 and MCP1, by abdominal subcutaneous and neck area adipocytes (27).

Adipocytes from the neck, especially the DN, area play a significant role in maintaining whole body energy homeostasis by performing continuous non-shivering thermogenesis (42), (43), (44), (45). However, the effect of irisin during the differentiation of SC and DN area adipocytes has not yet been elucidated. Recent publications pointed out that irisin may induce a different degree of browning response based on the origin of the human adipose tissue (21), (46). According to our results presented here, irisin did not directly influence the expression of thermogenesis-related genes in the SC and DN area adipocytes. However, it induced components of a secretory pathway leading to the release of *CXCL1*.

The targeted genetic impairment of the thermogenic capacity of BAT in mice (e.g. Ucp1^-/-^ mice) results in a less pronounced phenotype than the ablation of BAT (17). Transplantation of small amounts of BAT or activated beige adipocytes leads to significant effects on systemic metabolism, including increased glucose tolerance or attenuated fat accumulation in the liver in response to an obesogenic diet (47). Further studies highlighted the important secretory role of BAT, leading to an increased interest in identifying batokines in rodents that can exert autocrine, paracrine or endocrine effects. Several recently discovered batokines, such as FGF21, NRG4, BMP8b, CXCL14, or adiponectin have been shown to exert a protective role against obesity by enhancing beiging of WAT, lipolysis, sympathetic innervation, or polarization of M2 macrophages (16). We found that IL-6, released as a batokine, directly improves browning of human abdominal subcutaneous adipocytes (27). Our findings suggest that CXCL1 is a novel batokine, which can be secreted in response to specific cues. This is further supported by gene expression data from single cell analysis of human subcutaneous adipocytes; in thermogenic cells, genes of *CXCL1*, and other secreted factors, such as *CXCL2, CXCL3, CXCL5, CCL2*, and *IL6*, were significantly upregulated in response to forskolin that models adrenergic stimulation of heat production (48).

CXCL1 is a small peptide belonging to the CXC chemokine family. Upon binding to its receptor, CXCR2 (49), it acts as a chemoattractant of several immune cells, especially neutrophils (50). CXCL1 initiates the migration of immune and endothelial cells upon injury-mediated tissue repair (28). Conditioned medium containing CXCL1, collected during differentiation of SC and DN adipocytes in the presence of irisin, significantly improved the adhesion of HUVECs. We observed the similar response when they were directly treated with the recombinant chemokine (Figure 6D). Together this indicated a beneficial paracrine role of the released CXCL1 from differentiating adipocytes upon irisin treatment.

Our study shed light on an important role of irisin, as a regulator of batokine release from differentiating adipocytes of the neck area. The study also indicated the upregulation of various other cytokines, such as *CX3CL1, IL32, CXCL2, IL34, CXCL5*, and *CXCL3*. Further studies are required to reveal the impact of irisin stimulated release of other cytokines, which may have beneficial effects on local tissue homeostasis or metabolic parameters of the entire body.

Irisin can exert non-thermogenic effects on several tissues, including the liver (51), central nervous system (52), (53), blood vessels (54), or the heart (55). In mouse osteocytes, irisin acts via a subset of integrin receptor complexes, which are assembled from ITGAV and either ITGB1, ITGB3, or ITGB5 (20). These integrins transmit the effect of irisin in inguinal fat and osteoclasts *in vivo* (20), (56). In our experiments, RT-qPCR analysis of *ITGAV* expression has revealed its high expression in both preadipocytes and differentiated adipocytes, which was further upregulated upon irisin treatment (Figure 1D). RNA Sequencing also proved that the β-integrin subunits were abundantly expressed in both preadipocytes and differentiated adipocytes (Supplementary figure 1). However, RGDS peptide exerted only a moderate effect on the irisin-stimulated CXCL1 secretion. This suggests that irisin initiates some of its biological effects via other, currently unknown receptor(s) as well. The canonical integrin signaling includes the phosphorylation of FAK and Zyxin, followed by phosphorylation of AKT (at T308) and CREB (20). However, other studies proposed positive effects of irisin on cAMP-PKA-HSL (57), AMPK (58), (59), or p38 MAPK (18) pathways. Of note, RGDS peptide was applied at a relatively low concentration, in which anoikis was not observed. It is still possible that some of the administered irisin still access their integrin receptors at this condition.

It has already been reported that *CXCL1* gene expression is directly controlled by NFκB (60). NFκB-signaling might be induced in *ex vivo* differentiated adipocytes by released saturated fatty acids that can activate toll-like receptor (TLR) 4, which is abundantly expressed at mRNA level in hASCs and adipocytes of human neck (data not shown) (61), (62). Our data indicate that genes of canonical NFκB-signaling, which are abundantly expressed in neck area adipocytes, are upregulated when differentiated in the presence of irisin (Figure 5A,B). The absence of TNFα or IL-1ß-upregulation and release during the differentiation in the presence of irisin excluded the possibility of endotoxin contamination of the recombinant hormone. Although, irisin was reported previously to inhibit LPS-induced NFκB activation (63), (64), adipocytes differentiated in the presence of both SN50 and irisin released less CXCL1 than those of treated with irisin alone (Figure 5E). Further research is needed to explore the irisin-induced molecular events in the distinct human adipocyte subsets.

## 6 Author Contributions

LF, EK, AS and RK conceived and designed the experiments. AS, EK, SP, RK, and AV performed the experiments. EK, AS, and AV generated primary cell cultures for the experiments. BBT analysed the RNAseq data. AR analysed and visualized gene interaction networks. ICs, AS, AV, and ZsB performed microscopy and image analysis. FGy provided tissue samples, IRK-Sz provided HUVEC cells. AS and EK wrote the manuscript with inputs from BBT. LF mentored the writing and revised the draft. LF, EK, and IRK-Sz acquired funding. All authors approved the submitted version.

## 8 List of non-standard abbreviations

BAT: Brown adipose tissue
DN: Deep-neck derived adipocytes
GRO: Growth-related oncogene
hASCs: Human adipose-derived stromal cells
HUVEC: Human Umbilical Vein Endothelial Cells
IgG: Immunoglobulin G
IL: Interleukin
MCP1: Monocyte chemoattractant protein 1
NFκB: Nuclear factor-κB
PI: Propidium iodide
SC: Subcutaneous neck derived adipocytes
WAT: White adipose tissue

## 9 Acknowledgments

We thank Jennifer Nagy for technical assistance and Dr. Zsuzsa Szondy for reviewing the manuscript.

## 10 Ethics statement

The study protocol has been approved by Medical Research Council of Hungary (20571-2/2017/EKU). Experiments were performed strictly in accordance with the approved ethical regulations and guidelines.

## 12 Supplementary Material

## 13 Data availability statement

RNA-seq data was deposited to [Sequence Read Archive (SRA)] database [https://www.ncbi.nlm.nih.gov/sra] under accession number PRJNA607438. Other data that support the findings of this study are available from the corresponding authors upon reasonable request.

