## Supplementary Figures and Tables for "Irisin stimulates the release of CXCL1 from differentiating human subcutaneous and deep-neck derived adipocytes via upregulation of NFκB pathway"

### *Supplementary Material*

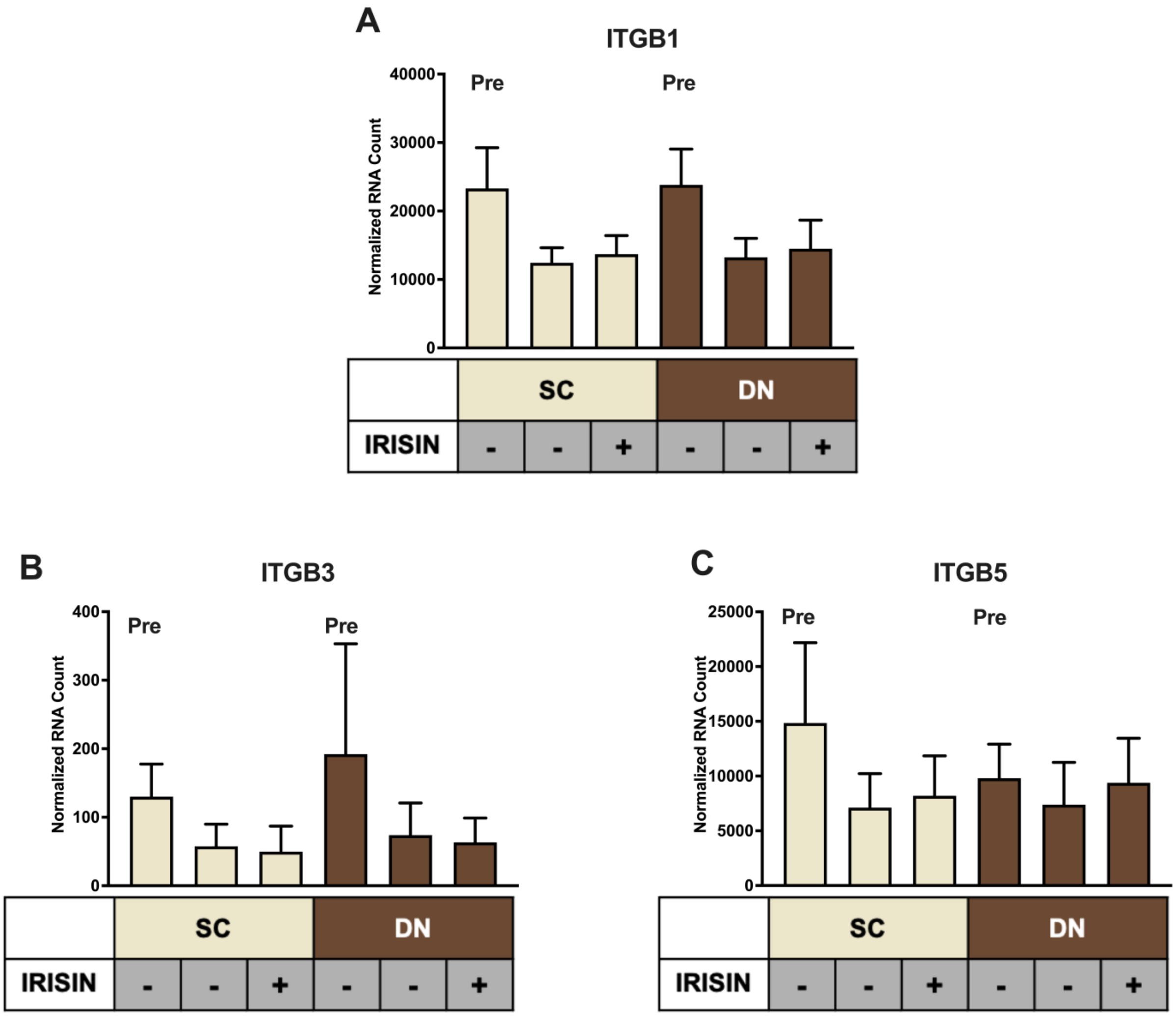


**Supplementary Figure 1. Proposed irisin receptor subunits are abundantly expressed in subcutaneous (SC) and deep-neck (DN) preadipocytes (Pre) and adipocytes of human neck.** SC and DN preadipocytes were differentiated and treated as in Figure 1. Quantification of gene expression of *ITGB1* (A), *ITGB3* (B), and *ITGB5* (C) by RNA-Sequencing (n=9). Data presented as Mean ± SD.

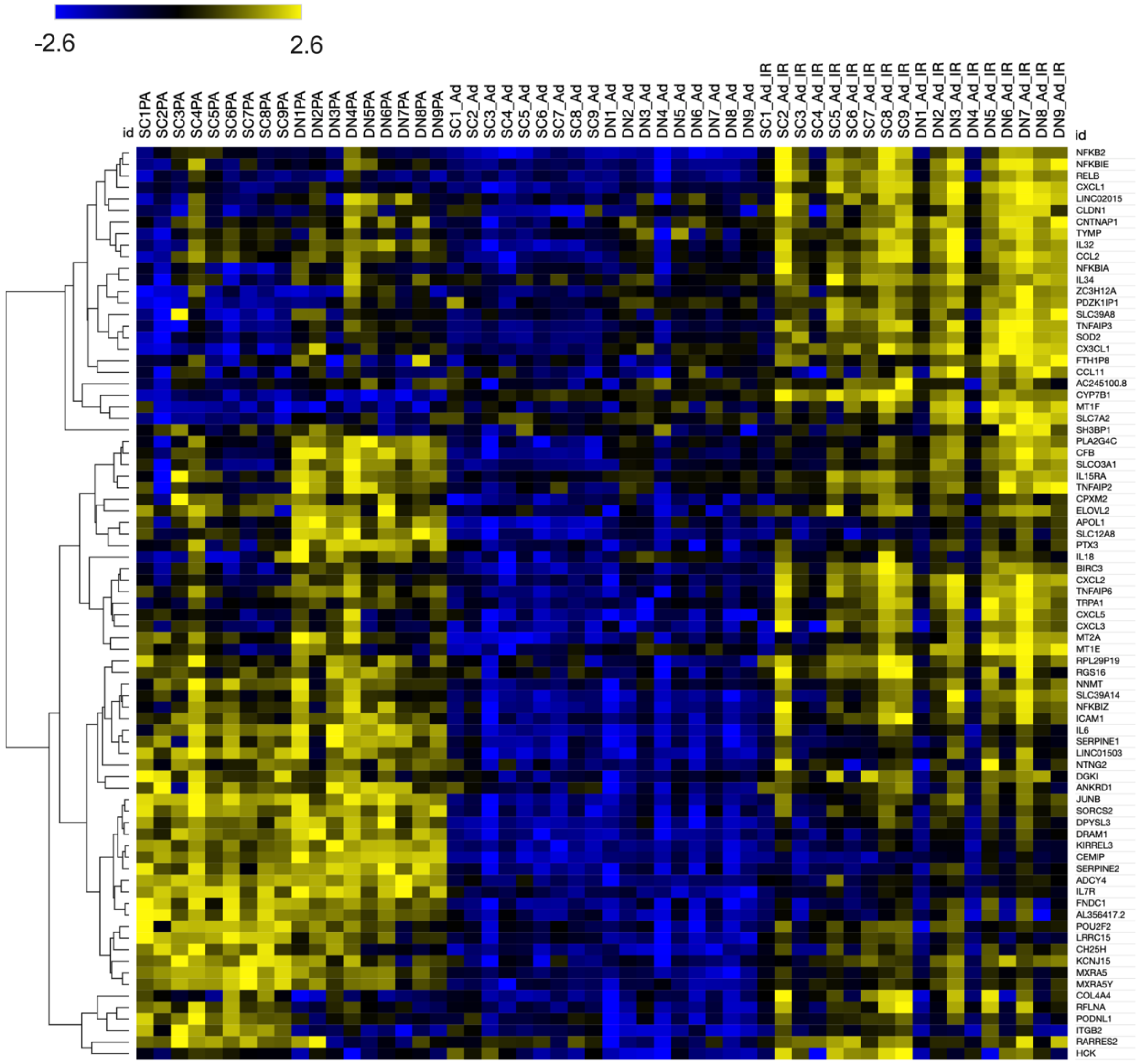


**Supplementary figure 2. Heatmap highlighting genes upregulated by irisin treatment in subcutaneous (SC) and deep-neck (DN) preadipocytes (Pre) and adipocytes (Ad) of human neck.** SC and DN preadipocytes were differentiated and treated as in Figure 1. Heatmap illustrating the expression of genes upregulated by irisin among all samples as evaluated by RNA Sequencing.

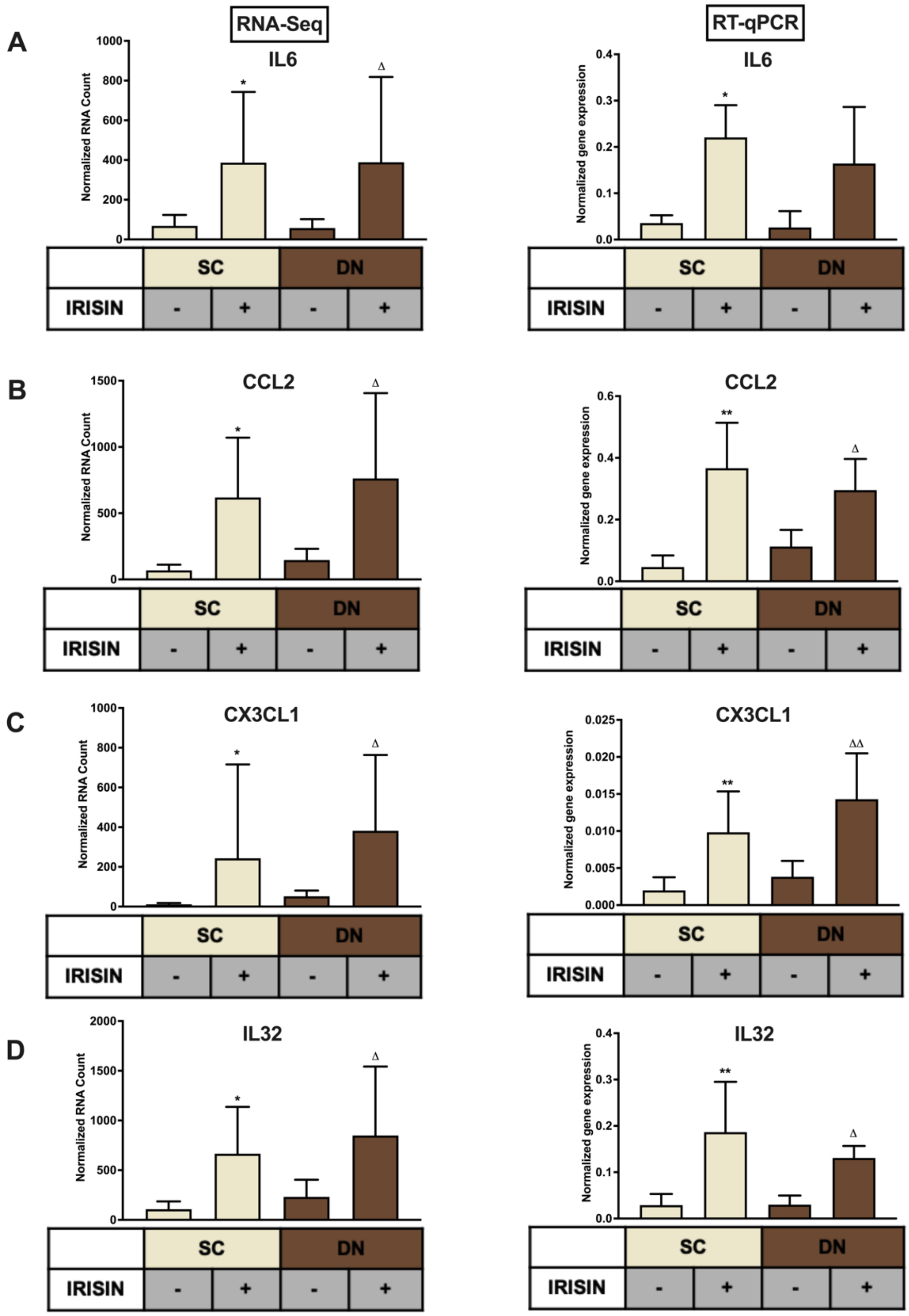


**Supplementary figure 3. Evaluation and validation of RNA Sequencing data for genes encoding cytokines, during the differentiation of subcutaneous (SC) and deep-neck (DN) derived adipocytes upon irisin treatment.** SC and DN preadipocytes were differentiated and treated as in Figure 1. Quantification of gene expression of *IL6* (A), *CCL2* (B), *CX3CL1* (C), and *IL32* (D) as assessed by RNA Sequencing (left, n=9) and RT-qPCR normalized to *GAPDH* (right, n=5). Data presented as Mean ± SD. * : Refers to compared with SC, ∆ : Refers to compared with DN. *,$^{\Delta}$ $p<0.05$ and **, $^{\Delta\Delta}$ $p<0.01$. Statistics: GLM (RNA-Sequencing) and One-way ANOVA with Tukey's post-test (RT-qPCR).

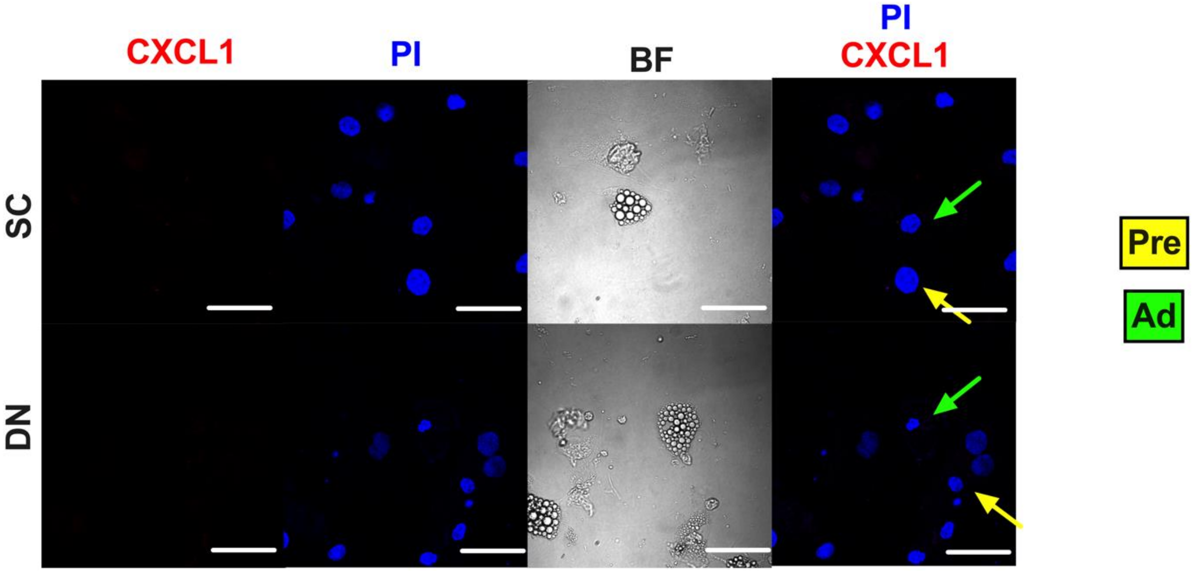


**Supplementary figure 4. Representative images of secondary antibody controls proving the specificity of CXCL1 immunostaining.** Scale bars represent 30 µm.

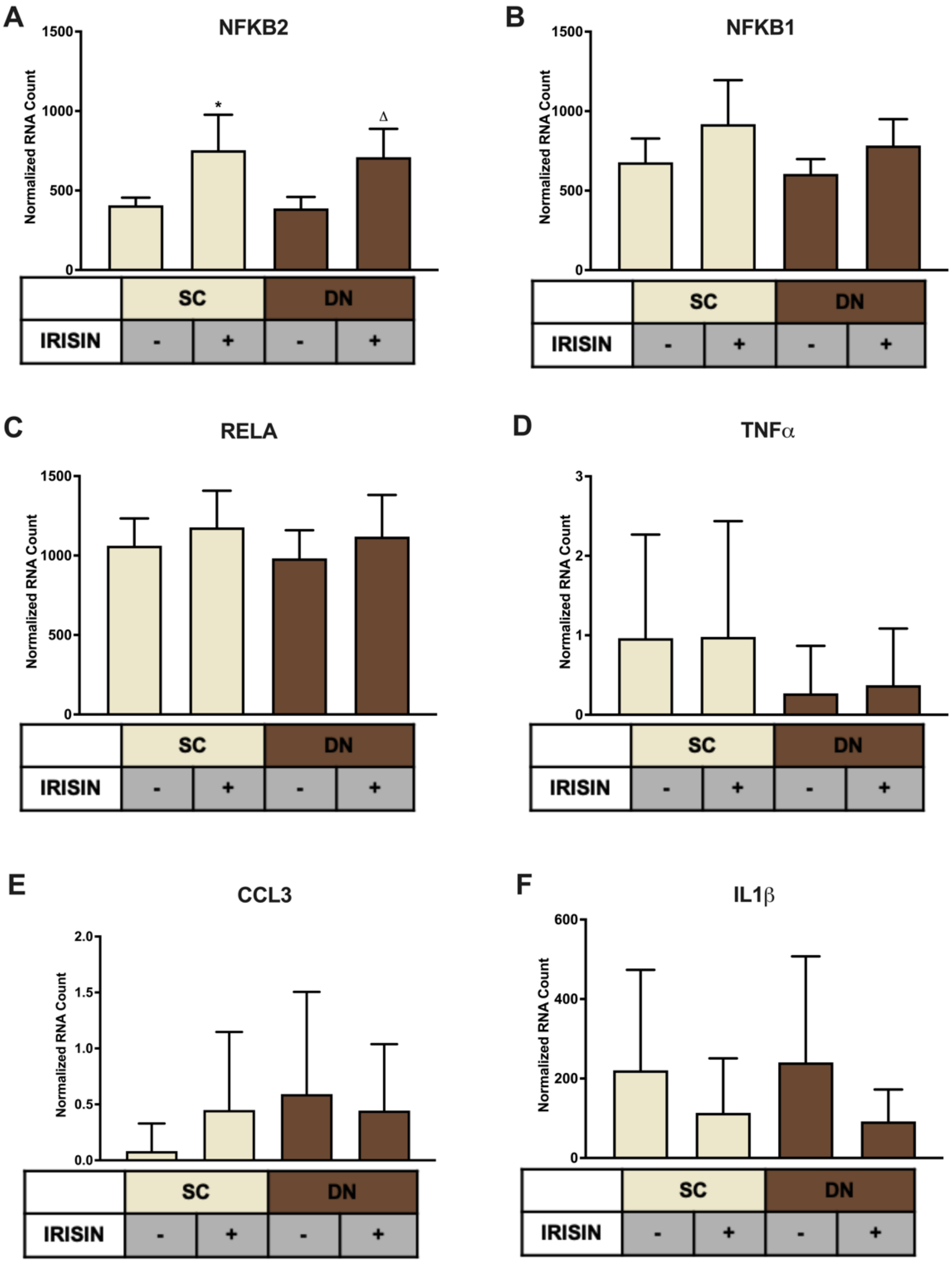


**Supplementary figure 5. Irisin treatment upregulated genes related to NFκB pathway during the differentiation of subcutaneous (SC) and deep-neck (DN) area adipocytes, while the expression of pro-inflammatory markers remained unchanged.** SC and DN preadipocytes were differentiated and treated as in Figures 1-4. Quantification of gene expression of *NFKB2* (A), *NFKB1* (B), *RELA* (C), *TNFα* (D), *CCL3* (E), and *IL1β* (F) as assessed by RNA Sequencing. Data presented as Mean ± SD. * : Refers to compared with SC, ▵ : Refers to compared with DN, *,▵ $p<0.05$, n=9 (Statistics: GLM)

**Supplementary Table 1. Significantly upregulated genes during the differentiation of subcutaneous (SC) and deep-neck (DN) derived adipocytes upon irisin treatment.**

| SC Irisin Upregulated | |
|---|---|
| **Gene-symbol** | **log2FoldChange** |
| *CXCL1* | 7.224 |
| *CXCL3* | 4.266 |
| *CXCL5* | 3.979 |
| *CX3CL1* | 3.813 |
| *CXCL2* | 3.684 |
| *TNFAIP6* | 3.581 |
| *CCL2* | 3.460 |
| *CFB* | 2.997 |
| *IL32* | 2.945 |
| *BIRC3* | 2.838 |
| *COL4A4* | 2.771 |
| *ICAM1* | 2.673 |
| *IL6* | 2.581 |

| DN Irisin Upregulated | |
|---|---|
| **Gene-symbol** | **log2FoldChange** |
| *CXCL1* | 6.258 |
| *CXCL5* | 5.154 |
| *CXCL2* | 3.970 |
| *CXCL3* | 3.387 |
| *HCK* | 3.374 |
| *TNFAIP6* | 3.073 |
| *CFB* | 2.926 |
| *SLC7A2* | 2.922 |
| *RFLNA* | 2.847 |
| *CLDN1* | 2.683 |
| *IL6* | 2.682 |
| *CCL11* | 2.664 |
| *CX3CL1* | 2.652 |

| | |
|---|---|
| *SOD2* | 2.478 |
| *CLDN1* | 2.349 |
| *TRPA1* | 2.345 |
| *RFLNA* | 2.327 |
| *LRRC15* | 2.272 |
| *ELOVL2* | 2.262 |
| *IL18* | 2.247 |
| *LINC02015* | 2.119 |
| *MT2A* | 2.002 |
| *RGS16* | 1.897 |
| *TNFAIP3* | 1.837 |
| *PDZK1IP1* | 1.818 |
| *CPXM2* | 1.794 |
| *AC245100.8* | 1.645 |
| *CYP7B1* | 1.579 |
| *POU2F2* | 1.572 |
| *SLC39A8* | 1.493 |

| | |
|---|---|
| *CCL2* | 2.465 |
| *ICAM1* | 2.460 |
| *TRPA1* | 2.411 |
| *MT1F* | 2.375 |
| *LRRC15* | 2.360 |
| *ELOVL2* | 2.282 |
| *SOD2* | 2.228 |
| *MT2A* | 2.220 |
| *AL356417.2* | 2.159 |
| *IL32* | 2.094 |
| *MXRA5Y* | 1.982 |
| *FNDC1* | 1.950 |
| *POU2F2* | 1.926 |
| *CH25H* | 1.902 |
| *ANKRD1* | 1.798 |
| *RGS16* | 1.757 |
| *TNFAIP3* | 1.735 |

| | |
|---|---|
| *KIRREL3* | 1.452 |
| *IL34* | 1.405 |
| *NFKBIZ* | 1.361 |
| *DPYSL3* | 1.324 |
| *SORCS2* | 1.302 |
| *APOL1* | 1.228 |
| *RELB* | 1.227 |
| *TYMP* | 1.187 |
| *RPL29P19* | 1.056 |
| *IL15RA* | 1.048 |
| *NFKBIA* | 1.042 |
| *NNMT* | 1.009 |
| *JUNB* | 0.999 |
| *DGKI* | 0.999 |
| *NFKBIE* | 0.995 |
| *CNTNAP1* | 0.908 |
| *ZC3H12A* | 0.901 |

| | |
|---|---|
| *IL7R* | 1.717 |
| *BIRC3* | 1.711 |
| *KCNJ15* | 1.691 |
| *SERPINE1* | 1.680 |
| *MXRA5* | 1.635 |
| *PDZK1IP1* | 1.604 |
| *NFKBIZ* | 1.583 |
| *KIRREL3* | 1.527 |
| *SLC39A8* | 1.523 |
| *SERPINE2* | 1.518 |
| *ITGB2* | 1.490 |
| *ADCY4* | 1.434 |
| *CEMIP* | 1.396 |
| *SLC39A14* | 1.374 |
| *LINC01503* | 1.336 |
| *PODNL1* | 1.332 |
| *PTX3* | 1.320 |

| | |
|---|---|
| *NFKB2* | 0.899 |
| *PLA2G4C* | 0.862 |
| *SLC39A14* | 0.849 |

| | |
|---|---|
| *IL34* | 1.317 |
| *RPL29P19* | 1.269 |
| *RELB* | 1.255 |
| *CYP7B1* | 1.166 |
| *NFKBIE* | 1.162 |
| *MT1E* | 1.150 |
| *NNMT* | 1.115 |
| *SH3BP1* | 1.097 |
| *JUNB* | 1.030 |
| *RARRES2* | 1.020 |
| *TNFAIP2* | 1.010 |
| *DRAM1* | 1.001 |
| *SLC12A8* | 0.986 |
| *SLCO3A1* | 0.977 |
| *NFKBIA* | 0.970 |
| *ZC3H12A* | 0.926 |
| *FTH1P8* | 0.916 |

| | |
|---|---|
| *NFKB2* | 0.902 |
| *NTNG2* | 0.888 |
